# How does immediate auditory feedback coupled to locomotion influence the spontaneous actions of rats?

**DOI:** 10.1101/2024.12.18.628882

**Authors:** Kosuke Yoshida, Reo Wada, Shiomi Hakataya, Genta Toya, Kazuo Okanoya, Hiroki Koda

## Abstract

Sense of agency is a subjective experience that is difficult to examine in animals and preverbal infants. In animals and infants, the sensory monitoring of self-motion, which is the basis of this sense, and the contingency between self-motion and sensory feedback are investigated. Here, we performed experiments on rats using locomotor movements and auditory feedback of pulsed sounds with real-time changes in pulse interval corresponding to the velocity of locomotion, and tested whether rats change their locomotion depending on the degree of contingency between locomotion and auditory feedback. The results showed that rats increased their locomotion distance when the pulse interval of sound changed in accordance with the velocity of locomotion. However, this effect was not long-lasting. This study provides a simple and convenient method to examine the awareness of the contingency between self-motion and auditory feedback in a wide range of animal species without long-term training, and is expected to be a useful tool for comparisons with humans.

## 1. Introduction

Do the animals recognize the sense that the animals themselves have influence their environment through their own actions? Such a sense is called sense of self-agency and is thought to be the basis for the emergence of self-knowledge and consciousness (Gallagher, 2000). However, since this sense is a subjective experience, it is difficult to measure in animals and pre-verbal infants. Experimental findings on the acquisition and evolution of self-agency, with respect to ontogeny and phylogeny, are still scarce (Terrace & Metcalfe, 2005).

One of the key features of this sense is the temporal contingency of the executed action and the immediate sensory feedback that occurs as a result of that action (Blakemore et al., 2002). Animals including humans continuously monitor this contingency of action and sensory feedback, comparing the intended action planning with the actual results of the action and evaluating the error. Focusing on this sensory feedback, sense of agency has been examined by observing behavioral changes produced by manipulating the degree of sensory feedback. For example, spontaneous changes in actions such as hand action or movement operation have been observed by adding temporal delays and spatial deviations to the sound and cursor display on the screen (e.g., Franck et al., 2001; Fourneret & Jeannerod, 1998; Synofzik et al., 2006). In developmental science, a study was conducted to examine infants’ behavior by connecting their limbs to an overhead mobile which provided visually changing stimuli when the infants moved their legs or arms (Rovee & Rovee, 1969). The results showed that the frequency of pulling increased when they were connected to the mobile above them. This is evidence of self-exploratory behavior, which is argued to be related to the emergence of self-agency. In a similar experiment, it is known that infants show an exploration-like sucking when given a pacifier which changes the feedback stimulus corresponding to the sucking pressure (Rochat & Striano, 1999). Eye-tracking experiments using an “image-scratching” paradigm, in which the visual features depicted on a monitor change depending on how long the infant gazes at the monitor, also show self-exploratory behavior (Miyazaki et al., 2014). The infants’ exploratory behavior increased when given immediate sensory feedback, suggesting the existence of a cognitive process that is the basis of self-agency.

Even in animals that do not verbally report their subjective experiences, the same method as in pre-verbal infants provides an effective means of understanding the phylogenesis of self-agency. Here we implemented an experimental apparatus in which auditory feedback is constantly played with or without the contingency between the velocity change of the rat and the sensory feedback (*n* = 30). If the rats recognize the contingency, we would expect self-exploratory behavior to be elevated in the movement-sound-contingent feedback condition, as is the finding in human infants. Specifically, we would expect an increase in total distance traveled and a transient increase in movement velocity. Lastly, the general applicability of the method to a wide range of animal species was examined using the techniques of animal tracking from video and acoustic feature changes of sound stimuli based on movement velocity.

## 2. Materials and methods

### Animals

We used 30 male Sprague-Dawley rats (Japan SLC, Inc., Shizuoka, Japan). All rats were 9-10 months old. They were kept in separate cages of 4 rats each, totally in 8 cages. Of all 32 animals, 30 were used for experiments and 2 for preliminary testing the apparatus. The housing environment was controlled with 12L:12D daylight cycle (lights off at 7:00 pm), room temperature between 20-26°C, and humidity between 40-60%. They were fed daily with a fixed amount (app. 31g/individual) of pellets and allowed to drink water freely.

### Animal tracking

We used a custom-made square open field chamber (800 mm W x 800 mm D x 900 mm H, Figure 1A) in which one rat could move freely. An infrared USB camera (See3CAM_CU27) was installed above the chamber to provide a bird’s-eye view of the chamber at all times. The experiment was conducted at night and the experimental room was kept in dim light. The video acquired by the camera was constantly captured on a laptop computer (mouse K7-I7G1BBK-A, CPU: intel Core i7, GPU: NVIDIA GeForce 1650, OS: Windows 11), with 640 x 480 pixels of 30 fps of AVI format. The rats were detected from the videos, and real-time tracking and judgment of their position on the plane and velocity of locomotion were performed. For real-time detection, we used Yolov8 and implemented our own code to determine the rat’s position. The detection model was retrained by transfer learning of the existing Yolov8 model with annotated supervised data that we prepared ourselves. The center of gravity was calculated from the detected rat’s bounding box and tracked as the rat’s position. The center-of-gravity position (*x_t_*, *y_t_*), where *x*, *y* are the vertical and horizontal positions on the movie and t is the step time to calculate) was calculated every 100 msec. From the time series data of the center of gravity position, the primary differential between the center of gravity position (*x*_*t*-1_, *y*_*t*-1_) at the previous computation step time and the current center of gravity position (*x_t_*, *y_t_*) was calculated, and the Euclidean distance was defined as the current velocity, i.e., 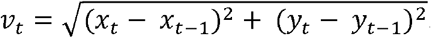

**Figure 1.**
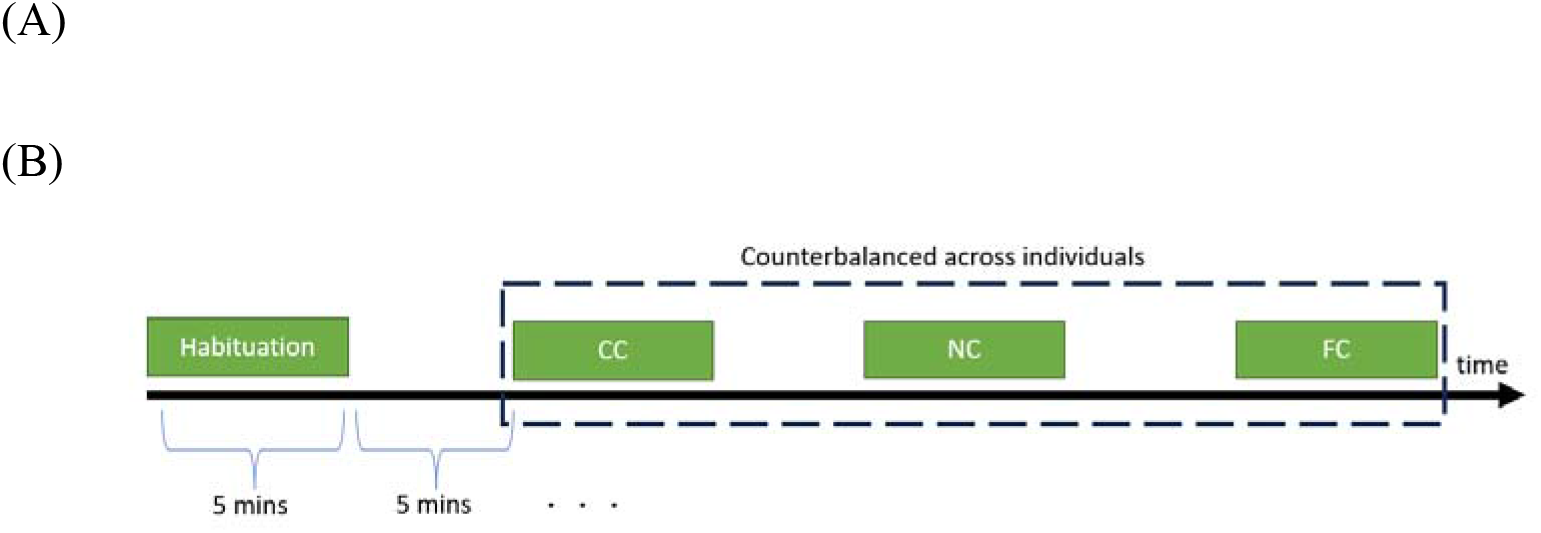
(A) Schematic illustration of the apparatus from the top view (left), and from the side view(right). The subject rat could move freely in the chamber, above which a speaker and a camera were installed. (B) Procedure of the experiment. One rat was placed in an open field chamber for 5 minutes without feedback sound (habituation phase). Following the habituation phase, we proceeded to the actual test phase, in which three auditory feedback conditions (FC, NC, CC) were set, each lasting 5 minutes. We randomized the order of the conditions and counterbalanced them across individuals, always inserting a 5-min break after one condition to complete all conditions.

### Auditory feedback

During the experiment, we set two tempos (10 Hz with 100 msec inter-stimulus interval or 2 Hz with 500 msec inter-stimulus interval) of 10000 Hz and 80 msec pure tone stimuli and continued to play them at either tempo. For the two tempos, we set three playback conditions. First, in the fixed pulse interval feedback condition (Fixed condition, FC), pure tone stimuli were always played back at a tempo of 2 Hz, independent of the rats’ velocity of locomotion. Next, in the non-contingent feedback condition (Non-contingent condition, NC), the pure tone stimulus was played independently of the rats’ velocity, as in the Fixed condition, but the tempo was randomly switched between 2 Hz and 10 Hz. Finally, in the movement-sound-contingent feedback condition (Contingent condition, CC), the tempo was switched with the rats’ velocity of locomotion according to the following criteria: 1) normally, the tempo was played at 2 Hz; 2) if the velocity exceeded 3-pixel values, the tempo was switched to 10 Hz; and 3) if the velocity was less than 3-pixel values, the tempo was switched immediately to 2 Hz. The rat’s position detection, calculation of velocity of movement, playback of the tempo-switching sound stimulus, and logging of the experiment were all coded independently using the python module pygame v2.0>.

### Procedures

The experiment consisted of two phases: a device habituation, and test phases. First, one rat was placed in an open field chamber for 5 minutes without feedback sound (device habituation phase). This is the habituation phase to the new device. Following the device habituation phase, we proceeded to the actual test phase, in which three auditory feedback conditions (FC, NC, CC) were set, each lasting 5 minutes. We randomized the order of the conditions and counterbalanced them across individuals, always inserting a 5-min break after one condition to complete all conditions (Figure 1B). During each break, the subject was removed from the apparatus and spent the break outside the apparatus. After a 5-min break had elapsed, the subject was placed back into the apparatus, and recording started 10 seconds later.

### Statistical analysis summary

In the statistical analysis, we examined our data from multiple perspectives, using two different methods. First, we determined and conducted the method of analysis prior to the experiment (here, formal analysis). For the analyzed data, we calculated the cumulative distance traveled per minute, and conducted a generalized linear mixed model (GLMM) to explain the cumulative distance traveled, and estimated parameters to examine the effects of the factors. Additionally, apart from the formal analysis, model selection procedure for the generalized linear model was conducted as a post hoc analysis to estimate the best model and to examine the parameters included in that model (see SI method for details).

## 3. Results

Figure 2 represents the cumulative distance traveled for each interval of the three conditions (CC, NC, FC), for each elapsed time of a 1-minute interval. The parameter coefficient of the NC of the “*condition*” was estimated to be negative (*β ± SE* = −176±79, *t* = −2.24, *p* = 0.026, intercept of the model was set to “NC” and “1-minute interval” of th*etime*, Table S3), while the parameter coefficient of FC remained the same at 0 (*β ± SE* = −95±79, *t* = −1.22, *p* = 0.22). Negative values were estimated for the “*time*” (*β ± SE* = −65.7±17, *t* = −3.92, *p* = 0.00010). The interaction effect terms (“*conidtion* * *time*”) were estimated as 0 (*β ± SE* = 36.9±24, 24.8±24, *t* = 1.55, 1.04, *p* = 0.12, 0.29 for NC:Time or FC:Time). The results show that rats traveled particularly actively in the first 1-min interval in all conditions, especially in the CC, where they traveled longer distances than in the NC.

**Figure 2.**
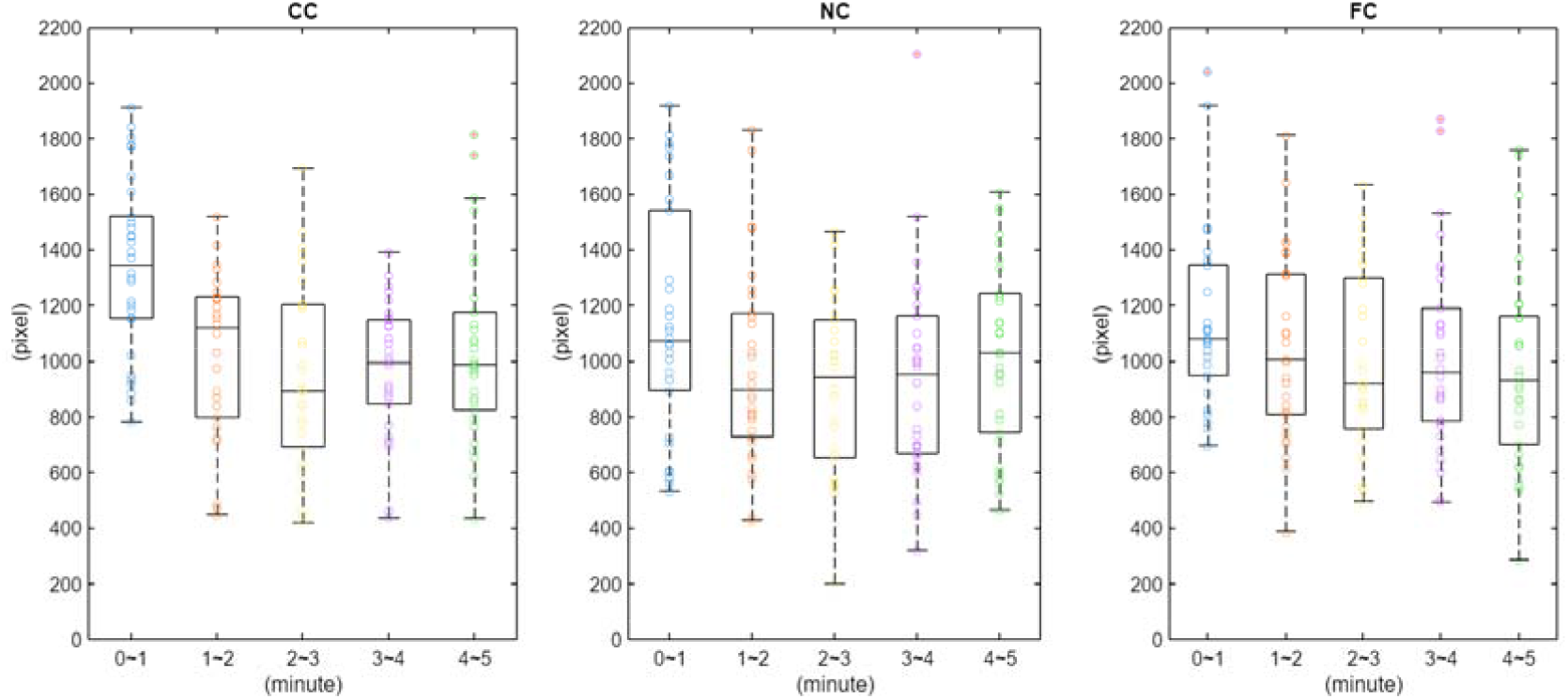
Boxplots of the cumulative distance traveled for each interval of three conditions. Each dot represents the distance traveled by each individual in each interval. The distances traveled by the rats decreased with time elapsed, and the distance traveled by the rats in the CC was longer than in the NC or FC, especially at the beginning of the experiment.

Since the visual inspection of the cumulative distance did not clearly confirm a linear decrease or increase according to time: therefore, we treated *time* parameter as a categorical variable in the post-hoc GLMM. When the model selection procedure was performed as a post-hoc analysis, results were mostly consistent: the smallest AIC model also included “*conidition*” and “*time*” and its interaction effect term (see Table S4 for AICs in model comparisons), supporting the consistent results of parameter coefficient estimations fitted by GLMM above. In addition, this model had a smaller AIC than the model considered in the formal analysis (see Table S3 and Table S4). However, the parameter coefficient of the FC was estimated to be negative (*β ± SE* = −168±72, *t* = −2.67, *p* = 0.02). In other words, this result suggests that 1) the distance traveled by the rats decreased with time elapsed, and 2) the distance traveled by the rats in the CC was longer than in the NC or FC, especially at the beginning of the experiment.

## 4. Discussions

The GLMM results indicated that the rats became more active and prolonged the movement trajectory distance immediately after the beginning of the experiment (the beginning of the test phase), when they were in the Contingent Condition (CC). This suggests that the rats could have become aware of the sound change in response to the movement change immediately after the introduction. On the other hand, this effect disappeared quickly. For this effect to be maintained or enhanced, sensory reinforcement, in which the change in the sound stimulus itself serves as a reinforcer, must be established. Sensory reinforcement is a phenomenon in which the behavior to detect the presented stimulus increases when the presented sensory stimulus itself becomes a reinforcer (Fujita & Matsuzawa, 1986). For example, the gazing time for “highly interesting” social stimuli such as the mother’s face or voice for babies, is spontaneously enhanced simply because of the baby’s motivation to view more of that stimulus. However, in general, in many animal studies, such social stimuli are less likely to be lasting rewards compared to food and water. Similarly, in the present experimental case of rats, this means that this self-exploration did not act as a sufficiently strong reward as a reinforcer. Therefore, we could not confirm the long-lasting effect of the self-exploration in rats.

The maintenance of self-exploratory behavior would be based on the awareness of the contingency. This association between awareness of contingency and maintenance of spontaneous self-exploratory behavior would be also related to the opposite aspect with learned helplessness, a well-known phenomenon in classical psychology of learning. Learned helplessness is a phenomenon investigated mainly by Seligman in animal experiments in which long-term exposure to uncontrollable aversive stimuli leads to the extinction of the escape behavior itself (e.g., Maier & Seligman, 2016; Overmier & Seligman, 1967; Seligman & Maier, 1967). Specifically, in an experimental box in which electric shocks are randomly generated, animals are exposed to conditions in which they can avoid danger by their own actions, such as pressing a lever, or conditions in which they cannot avoid danger by their own actions; the animals are then observed for their escape behaviors. Despite the same frequency of exposure to electric shocks, the animals, which cannot control their own escapes by their own actions, will not engage in spontaneous escape behavior or will take more trials leading up to the behavior. This is an example in which the occurrence of spontaneous behavior is inhibited by the violation of the contingency between the animal’s own actions and the responses that result from those own actions. The fact that the disruption of contingency causes the disappearances of spontaneous behavior and, conversely, the promotion of contingency in our experiment increases spontaneous exploratory behavior, would suggest a strong coupling between contingency and the emergence of spontaneous behavior. Interestingly, learned helplessness is a common phenomenon in animals that undergo conditioning, reflecting a biological principle with deep phylogenetic origins, as demonstrated in fish (Padilla et al., 1970), sheep (Greiveldinger et al., 2009), and humans (Thornton & Jacobs, 1971). Likewise, the awareness of the contingency between own actions and sensory feedback, as well as the perception of contingency discrepancy, will also occur in many animals as the precursors to a sense of agency. Humans may be unique compared to other animals in that they repeatedly self-explore and confirm the contingency of their own motor and sensory feedback.

Sense of self-agency is difficult to measure directly in animals because it is a subjective experience. However, the basis of sense, i.e., monitoring of own actions by motor manipulation and immediate sensory feedback, and automatic motor adjustment, is a universal sensorimotor system in all animals. In non-human primates such as chimpanzees and macaques, which show the remarkable motor control of hand or fingers, psychological experiments were conducted to manipulate the contingency between voluntary motor control and visual sensory feedback, and to test the presence or absence of sense of agency. For example, experiments using a trackball to judge the sense of agency for cursor manipulation were conducted, and the degree of spatiotemporal contingency discrepancy affected the judgments, suggesting the precursor of presence of self-agency in animals (Kaneko & Tomonaga, 2011). On the other hand, this method is not suitable for non-primate animals because it requires a long-term training, is so difficult that all individuals cannot be successfully trained, and is mostly limited to hand movements and visual feedback. The present experimental method uses auditory stimuli and locomotion in space, and is applicable to almost all animals, making it highly general-purpose. Auditory stimuli can provide feedback over a wide area through a loudspeaker, and locomotion in space can be easily evaluated in all animals, independent of specific actions such as hand movements. In particular, nocturnally adapted rats would be more suited to auditory feedback and body locomotion than visual stimulus feedback and specific body actions, because of their limited ability to perform visual tasks by body actions. The flexibility to change feedback methods, such as delayed feedback or motion-dependent pitch changes, would also be beneficial for comparative studies with a wide range of animal species. In terms of versatility and ease of implementation in general animals, our method contributes to the development of a new paradigm, namely, that a simple integration of animal locomotion and auditory feedback is all that is required.

## Supporting information

Supplemental method and result tables

## Acknowledgements

We appreciate Noriko Kondo, Sota Kikuchi, Sana Kohmoto for supporting daily care of animals. The study was supported by the Japan Society for the Promotion of Science (Grant □ in □ Aid for Scientific Research (S, #23H05428) to KO, (A, #21H04421; B, # 22H03914) to HK, (# JSPS fellow, # 21J23113) to SH, and by World-leading Innovative Graduate Study Program of Advanced Basic Science Course (WINGS-ABC) to RW.

