## Supplemental method and result tables for "How does immediate auditory feedback coupled to locomotion influence the spontaneous actions of rats?"

Supplementary materials for “How does immediate auditory feedback coupled to action influence the spontaneous actions of rats?”

Kosuke Yoshida^1^, Reo Wada^1^ , Shiomi Hakataya^13ab^, Genta Toya^23^, Kazuo Okanoya^13^, Hiroki Koda^14^

Supplementary Information for Methods (SI Methods)

**Analysis details**

*Formal analysis*

The following analysis procedures were determined prior to the experiment. We calculated the cumulative distance traveled per minute and considered its temporal changes as the target of our analysis. For the cumulative distance, we constructed a generalized linear mixed model (GLMM) with the following model formula,

$$distance \sim condition * time+ (1|individual)$$

where $condition$ is the auditory feedback condition (FC, NC, CC), $time$ is the time elapsed for each minute interval (integer from 1 to 5), and the parentheses represent random effects, with individual differences designated as intercepts for random effects. The results were estimated for the parameters and the probability of deviation from 0. For GLMM, Matlab's fitglme was used.

*Post hoc analysis*

A model selection procedure was also performed to compute the "best" model, i.e., the minimum AIC model, by means of a stepwise model selection procedure. For the model selection procedures, we used `dredge` function in `MuMIN` package with the GLMM fitted by `lmer` function of `lme4` in R.

Supplementary Tables

Table S1. The parameter coefficients of GLMM in the formal analysis

| Parameter | Estimate | SE | tStat | df | pValue |
| --- | --- | --- | --- | --- | --- |
| Intercept | 1260.3 | 64.4 | 19.6 | 444 | 5.23e-62 |
| NC | -176.2 | 78.6 | -2.24 | 444 | 0.026 |
| FC | -96.0 | 78.6 | -1.22 | 444 | 0.22 |
| Time | -65.7 | 16.8 | -3.92 | 444 | 1.03e-4 |
| NC:Time | 36.8 | 23.7 | 1.55 | 444 | 0.12 |
| FC:Time | 24.8 | 23.7 | 1.04 | 444 | 0.30 |

Table S2. The parameter coefficients of GLMM in the posthoc analysis.

i,v denotes “interval”.

| Parameter | Estimate | SE | tStat | df | pValue |
| --- | --- | --- | --- | --- | --- |
| Intercept | 1338 | 60.5 | 22.1 | 435 | 3.67e-73 |
| NC | -192.5 | 72.0 | -2.67 | 435 | 0.0078 |
| FC | -168.2 | 72.0 | -2.33 | 435 | 0.020 |
| Second i.v | -308.7 | 72.0 | -4.28 | 435 | 2.25e-05 |
| Third i,v | -396.6 | 72.0 | -5.51 | 435 | 6.32e-08 |
| Fourth i.v | -368.7 | 72.0 | -5.11 | 435 | 4.66e-07 |
| Fifth i.v | -298.4 | 72.0 | -4.14 | 435 | 4.13e-05 |
| NC:Second | 140.8 | 101.9 | 1.38 | 435 | 0.17 |
| FC:Second | 179.9 | 101.9 | 1.77 | 435 | 0.078 |
| NC:Third | 160.7 | 101.9 | 1.58 | 435 | 0.11 |
| FC:Third | 228.6 | 101.9 | 2.24 | 435 | 0.025 |
| NC:Fourth | 156.1 | 101.9 | 1.53 | 435 | 0.13 |
| FC:Fourth | 220.1 | 101.9 | 2.17 | 435 | 0.031 |
| NC:Fifth | 176.4 | 101.9 | 1.73 | 435 | 0.084 |
| FC:Fifth | 103.5 | 101.9 | 1.02 | 435 | 0.31 |

Table S3. Model comparison table sorted by AIC. These models treat “time” as a continuous variable. Plus(+) represents the term included in the model.

| Model order | $condition$ | $time$ | $condition:time$ | df | AIC | ΔAIC |
| --- | --- | --- | --- | --- | --- | --- |
| 1 | + | -40.91 | + | 444 | 6367.1 |  |
| 2 | + | -36.89 |  | 446 | 6379.3 | 12.23 |
| 3 |  | -36.89 |  | 448 | 6395.1 | 28.01 |
| 4 | + |  |  | 447 | 6399.4 | 32.37 |
| 5 |  |  |  | 449 | 6415.2 | 48.14 |

Table S4. Model comparison table sorted by AIC. These models treat “time” as a categorical variable. Plus(+) represents the term included in the model.

| Model order | $condition$ | $time$ | $conditiom:time$ | df | AIC | ΔAIC |
| --- | --- | --- | --- | --- | --- | --- |
| 1 | + | + | + | 435 | 6290.5 |  |
| 2 | + | + |  | 443 | 6363.8 | 78.30 |
| 3 |  | + |  | 445 | 6386.3 | 95.82 |
| 4 | + |  |  | 447 | 6444.7 | 154.2 |
| 5 |  |  |  | 449 | 6462.0 | 171.54 |
